# Zn^2+^ triggered two-step mechanism of CLIC1 membrane insertion and activation into chloride channels

**DOI:** 10.1101/2021.10.25.465729

**Authors:** Lorena Varela, Alex C. Hendry, Joseph Cassar, Ruben Martin-Escolano, Diego Cantoni, John C Edwards, Vahitha Abdul-Salam, Jose L. Ortega-Roldan

## Abstract

The CLIC protein family displays the unique feature of altering its structure from a soluble form to a membrane-bound chloride channel. CLIC1, a member of this family, is found in the cytoplasm or in internal and the plasma membranes, with membrane relocalisation linked to endothelial disfunction, tumour proliferation and metastasis. The molecular switch promoting CLIC1 activation remains unclear. Here, cellular chloride efflux assays and immunofluorescence microscopy studies have identified Zn^2+^ intracellular release as the trigger for CLIC1 activation and membrane relocalisation. Biophysical assays confirmed specific binding to Zn^2+^, inducing membrane association and enhancing chloride efflux in a pH dependent manner. Together, our results identify a two-step mechanism with Zn^2+^ binding as the molecular switch promoting CLIC1 membrane insertion, followed by pH activation of chloride efflux.

## INTRODUCTION

The Chloride Intracellular Channel (CLIC) family consists of a group of highly homologous human proteins with a striking feature, their ability to change their structure upon activation from a soluble form into a membrane bound chloride channel, translocating from the cytoplasm to intracellular membranes (Tulk et al., 2000; Valenzuela et al., 1997). CLIC1 is the best characterised of the CLIC protein family. It is expressed intracellularly in a variety of cell types, being especially abundant in heart and skeletal muscle (Valenzuela et al., 1997). CLIC1’s integral membrane form has been found to be localised mostly in the nuclear membrane, although it is present in the membranes of other organelles and transiently in the plasma membrane. It has also been shown to function as an active chloride channel in phospholipid vesicles when expressed and purified from bacteria, showing clear single channel properties (Tulk et al., 2000, 2002).

CLIC1 has been implicated in the regulation of cell volume, electrical excitability (Averaimo et al., 2014), differentiation (Wang et al., 2011a), cell cycle (Valenzuela et al., 2000) and cell growth and proliferation (Tung and Kitajewski, 2010). High CLIC1 expression has been reported in a range of malignant tumours (Gritti et al., 2014; Setti et al., 2015; Tian et al., 2014; Wang et al., 2011b; Zhang et al., 2015; Zhao et al., 2015) and cardiovascular diseases such as pulmonary hypertension (Abdul-Salam et al., 2010) and ischaemic cardiomyopathy (Gronich et al., 2010). One common effect of CLIC1 in these disease mechanisms is its effect on endothelial function (Gurski et al., 2015; Xu et al., 2016). CLIC1 compromises endothelial barrier function and mediates cell growth, angiogenesis and migration (Knowles et al., 2011; Tung and Kitajewski, 2010). These vascular malformations have been linked to electrophysiological alterations in the cells caused, at least in part, by the CLIC1 chloride channel activity (Gurski et al., 2015). The activity and deleterious function of CLIC1 is modulated by its equilibrium between the soluble cytosolic form and its membrane bound form. Only CLIC1 in its channel form has been shown to have pro proliferative and angiogenic activity, and specific inhibition of CLIC1 channel halts endothelial injury, tumour angiogenesis and progression and vascular inflammation.

Despite its clinical importance, to date very little and conflicting information is available for CLIC1 membrane insertion mechanism, and the structure of the channel form is unknown. In-vitro oxidation with hydrogen peroxide causes a conformational change due to the formation of a disulphide bond between Cys24 and the non-conserved Cys59, exposing a hydrophobic patch that promotes the formation of a dimer (Littler et al., 2004), in a process that has been proposed to lead to membrane insertion (Goodchild et al., 2009). However, numerous studies have shown that oxidation does not promote membrane insertion (Singh and Ashley, 2006); with evidence pointing at pH (Fanucchi et al., 2008; Stoychev et al., 2009) or cholesterol (Hossain et al., 2016) as the likely activation factors. Thus, long standing inconsistencies in the data surrounding the molecular switch that transforms CLIC1 from its soluble form into a membrane bound channel has prevented further advances in the understanding of CLIC1 function.

In this study, we have explored the membrane insertion activation mechanism of CLIC1. We have discovered the activation of CLIC1 chloride efflux by intracellular Zn^2+^ in Glioblastoma cells, which also causes CLIC1 relocalisation to the plasma membrane. Finally, in-vitro studies with purified CLIC1 have shown Zn^2+^ driven activation of chloride efflux, membrane association, as well as CLIC1 specific binding to both Zn^2+^ and Ca^2+^.

## RESULTS

Divalent cation binding, specifically Ca^2+^, is involved in the membrane interactions of other protein families (Dubé et al., 2016; Rosengarth and Luecke, 2003; Sciacca et al., 2013). Interestingly, the CLIC *C. Elegans* and *Drosophila* homologs *exc*-4 and *Dm*CLIC1 both contain a calcium ion bound trapped during the expression and purification process. We therefore sought to understand if divalent cations may have a role in CLIC1 activation as a chloride channel.

The effect of divalent cation-driven membrane localisation on the chloride efflux activity of CLIC1 was tested in U87G cells using MQAE as a fluorescent reporter of the intracellular concentration of Cl^-^. With no treatment, the chloride efflux activity is only modestly inhibited by the CLIC inhibitor IAA-94 or the Zn chelator TPEN, and thus is likely not CLIC mediated. Ionomycin, an ionophore shown to increase both Ca^2+^ and Zn^2+^ intracellular concentrations, induced a significant increase (p=0.0123) of the Cl-efflux, as seen by the increase in MQAE fluorescence, that was reversed upon treatment with the Zinc chelator TPEN or with the CLIC inhibitor IAA-94 (p<0.005) (Xu et al., 2016) (Figure 1). Interestingly, TPEN inhibits the chloride conductance of the control cells to a level similar to IAA-94, suggesting the involvement of Zn^2+^ in the activation of CLIC1 chloride efflux activation.

**Figure 1.**
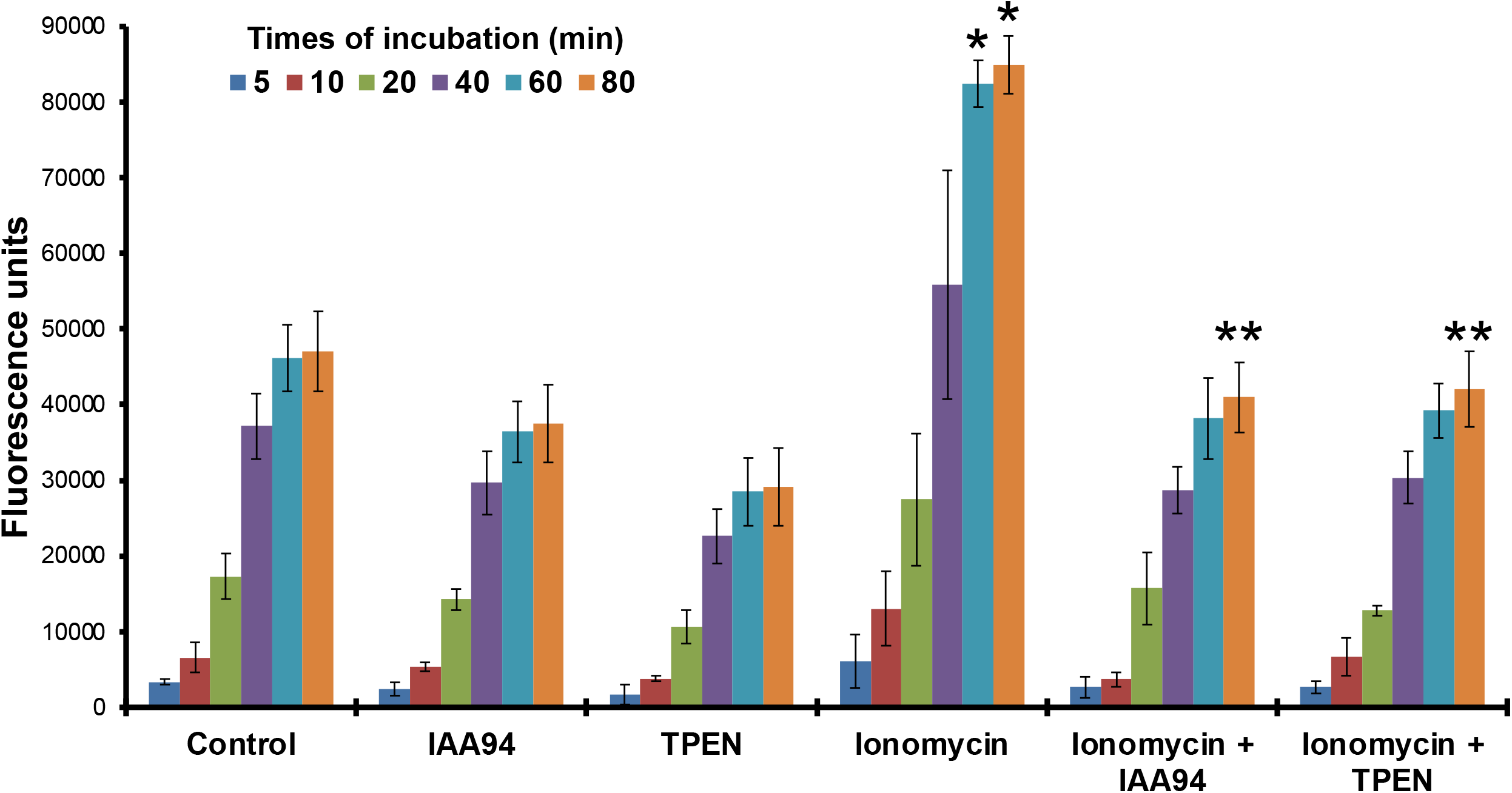
Effect of different treatments on chloride efflux (measured as fluorescence intensity units of the dye MQAE) in MQAE-stained U87 cells exposed to IAA94 (10 µM), TPEN (5 µM) and/Ionomycin (10 µM) for 80 min. Values constitute means of six independent determinations ± standard deviation. Ionomycin treatment resulted in statistically significant differences from 60 min compared to the control samples (p=0.0123) (* P, 0.05) and to all other treatments (** p<0.005) (n=6)

In light of these results, we questioned if Zn^2+^ could trigger CLIC1 membrane relocalisation in cells. Endogenous CLIC1 localisation was monitored in Human Umbilical Vein Endothelial Cells (HUVEC) and Glioblastoma (U87G) using immunostaining. CLIC1 typically exists in the cytosol in untreated HUVEC cells, and the addition of external ZnCl_2_ promoted the presence of CLIC1 at the plasma membrane. This effect was reversed with the treatment of TPEN or the chloride channel inhibitor NPPB (Figure 2A-D). A similar effect was observed in U87G cells. Addition of Ionomycin increased the degree of CLIC1 plasma membrane localisation (Figures E-G), and this effect was reversed by the addition of TPEN, in line with our findings in HUVEC cells and confirming that Zn^2+^ triggers the activation and membrane relocalisation of CLIC1.

**Figure 2:**
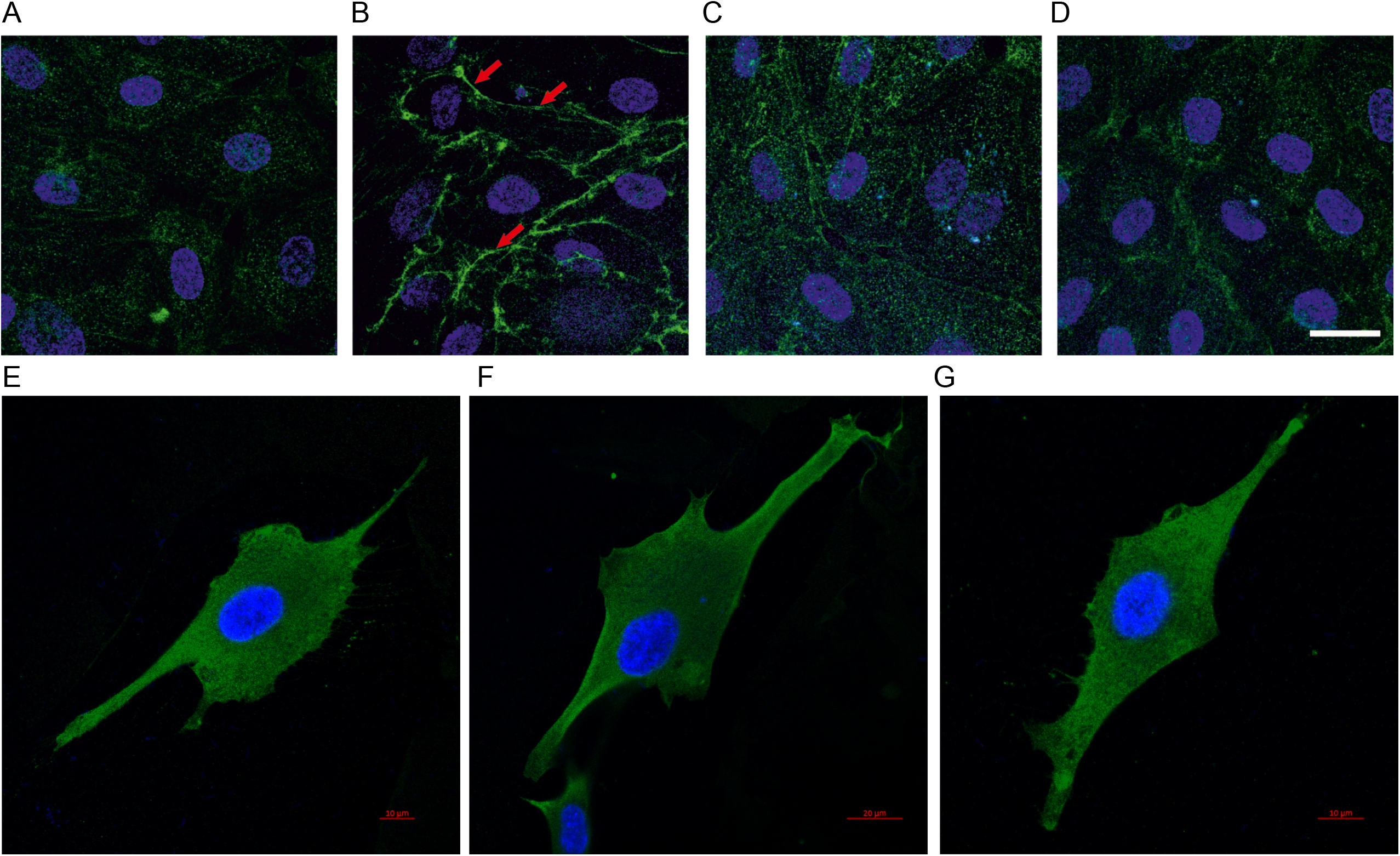
Divalent cations promote membrane insertion. **A-D** - CLIC1 localisation in Human Umbilical Vein Endothelial Cells (HUVEC). HUVECs were either untreated (A) or treated for 3 hours with (B) 10µM ZnCl2, (C) 10µM ZnCl2 with 5µM N,N,N’,N-tetrakis(2-pyridylmethyl) ethylenediamine (TPEN) and (D) 10µM with 10µM 5-nitro-2-(3-phenylpropyl-amino) benzoic acid (NPPB). CLIC1 typically exists in the cytosol and the addition of ZnCl_2_ promoted the presence of CLIC1 at plasma membrane (shown in red arrows). This effect was reversed with the treatment a zinc chelator (TPEN) or a chloride channel inhibitor (NPPB). **E-G** - Fluorescence microscopy images of U87G cells. U87Gs were either untreated (E), treated with 10 μM Ionomycin (F), or treated with 10 μM Ionomycin and x mM TPEN. Samples were stained with a CLIC1 antibody (green) and DAPI (blue).

Previous studies have indicated the importance of low pH for CLIC1 chloride. We used the previously reported vesicle-based chloride efflux assay to assess effect of either Calcium or Zinc ions on CLIC1 chloride channel activity in vitro (Tulk et al., 2002). The CLIC1 chloride channel shows strong pH dependence, with little activity above pH 6.5 (Warton et al., 2002). We measured channel activity at pH 5.5 and pH 7.5 in the nominal absence of divalent cations, or in the presence of 1μM Ca^2+^ or Zn^2+^ (figure 3A), using vesicles from the soybean lipid extract asolectin. The divalent cation concentration was chosen to be equimolar with the protein concentration in the reaction mix. Protein was mixed with cation prior to addition of vesicles. We found virtually no activity at pH 7.5 in the presence or absence of divalent cation. At pH 5.5, we found significant chloride permeability conferred by CLIC1 that was enhanced about 25% by the presence of Zn^2+^ (p=0.024), but not by the presence of Ca^2+^. We repeated the experiment using purified phospholipid vesicles rather than asolectin because of concern that the activity in the nominal absence of Zn^2+^ may be due to small amounts of Zn^2+^ contaminating the lipids. With purified phospholipids, the activity in the absence of Zn^2+^ was 35% lower than with asolectin vesicles, but the activation with 1 μM Zn was greater and amounted to a statistically significant 40% increase (p=0.0044).

**Figure 3.**
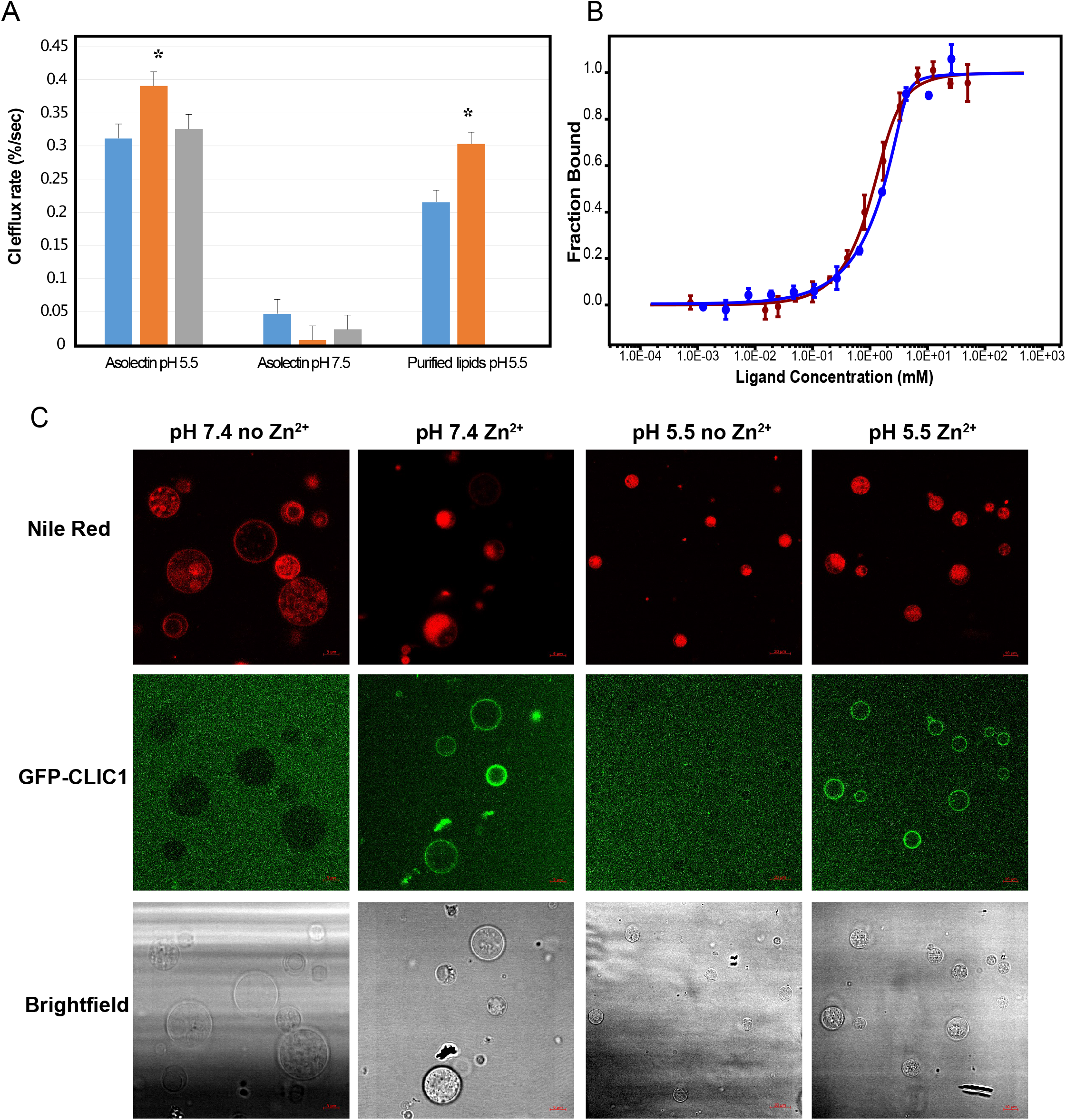
**A**- Zn^2+^ dependence on CLIC1 chloride efflux rate. Chloride efflux rates of CLIC1 in the absence of divalent cations (blue) or in the presence of stochiometric amounts of Zn^2+^ (orange) or Ca^2+^ (grey). Errors are shown as standard deviation (n=4). Highlighted with stars, Zn^2+^treatment resulted in statistically significant differences compared to the control samples in asolectin vesicles and using purified lipids at pH 5.5 (p<0.05, n=4) **B** – MST binding curves of CLIC1 titrations with Ca^2+^ (red) or Zn^2+^ (blue), displaying a Kd of 0.3mM ± 0.1mM for Ca^2+^ and 0.7 mM ± 0.2mM for Zn^2+^. Error bars represent the standard deviation. **C-** Fluorescent microscopy images of Asolectin. GUVs labelled with Nile red dye incubated with GFP-labeled CLIC1 in giant unilamellar vesicles at pH 7.4 (columns 1 and 2) and pH 5.5 (columns 3 and 4) in the absence (Columns 1 and 3) and presence (Columns 2 and 4) of Zn^2+^. Row 1 displays Red Nile fluorescence, row 2 CLIC1-GFP fluorescence and row 3 shows the brightfield images of the GUVs.

We subsequently attempted to separate activation of the CLIC1 chloride channel efflux activity from its insertion in the membrane. To this end, we explored the effect of Zn^2+^ and pH on membrane association of recombinantly expressed and purified GFP-tagged CLIC1 by incubating CLIC1-GFP with and without Zn^2+^ at pH 7.4 and 5.5 with giant unilamellar vesicles (GUVs). We used the fluorescence of Red Nile dye, which binds to the GUVs lipids, and GFP, to investigate the degree of co-localisation of CLIC1-GFP with the GUVs. Fluorescence microscopy images collected showed CLIC1 co-localisation in the membrane only in the presence of Zn^2+^ irrespective of the pH value, indicating that Zn^2+^ has a direct effect on CLIC1 triggering its membrane insertion (Figure 3C). Calcium on the other hand only induced partial insertion in the membrane at the same protein and cation concentrations, in agreement with its neglectable effect on CLIC1 chloride efflux (Figure S1).

Microscale Thermophoresis (MST) can be used to identify specific divalent cation binding on CLIC1 and to determine the affinity of these interactions. We used a C-terminal GFP tagged CLIC1 construct at a concentration of 1μM and at pH 7.4 to compare the binding affinities of Ca^2+^ and Zn^2+^ ions in the presence of asolectin vesicles, as CLIC1 can aggregate at molar ratios of Ca^2+^ and Zn^2+^ higher than 1 in the absence of lipids. CLIC1 showed specific binding to both ions, with Zn^2+^ and Ca^2+^having a similar apparent affinity of 300 μM (Figure 3B).

## DISCUSSION

### A model for CLIC1 membrane insertion

In this study, we demonstrate that CLIC1 binds to Zn^2+^, promoting membrane insertion in both model membranes and in a cellular environment. Once inserted, chloride efflux is only activated at low pH values. Combining our data, we propose a new two-step activation mechanism of CLIC1 whereby CLIC1 exists in a soluble state, and upon intracellular Zn^2+^ release CLIC1 binds to it, alters its structure and inserts into the membrane (Figure 4). The channel is only activated at low pH, suggesting either a pH gating mechanism or a conformational rearrangement within the membrane at low pH values, since pH had no effect on CLIC1 interaction with the membrane. CLIC1 has been shown to regulate phagosomal acidification (Salao et al., 2016), supporting the role of pH in the activation of the channel. An increase in the cellular Zn^2+^ concentration has been linked upstream and downstream of the ROS signalling pathway (Görlach et al., 2015), explaining previous studies on the relationship between CLIC activity with ROS production and oxidative stress. Ionomycin is known to increase both intracellular levels of Zn^2+^ and Ca^2+^. We show that although ionomycin treatment could result in CLIC1 binding to both Zn^2+^ and Ca^2+^ with similar affinity, only Zn^2+^ is able to induce membrane association (Figure S1) and chloride efflux (Figure 3A). This is further supported by treatment with the zinc chelator TPEN, that is able to suppress CLIC1 membrane relocalisation and its chloride conductance. The higher activation potential of Zn^2+^ over Ca^2+^ must therefore be a result of a higher degree of structural changes resulting from the interaction with Zn^2+^. The apparent affinity of CLIC1 towards Zn^2+^ and Ca^2+^ determined by MST is weak, while at molar ratios of 1:1 Zn^2+^ significantly increases CLIC1chloride conductance. MST relies on the presence of a fluorescent molecule attached to the protein, and in this case, we used a GFP fused to the C-terminus of CLIC1. We hypothesise that the presence of this C-terminal GFP may be interfering with the structural changes that need to occur for CLIC1 to bind to Zn^2+^and to insert in the membrane, lowering the affinity for this interaction.

**Figure 4.**
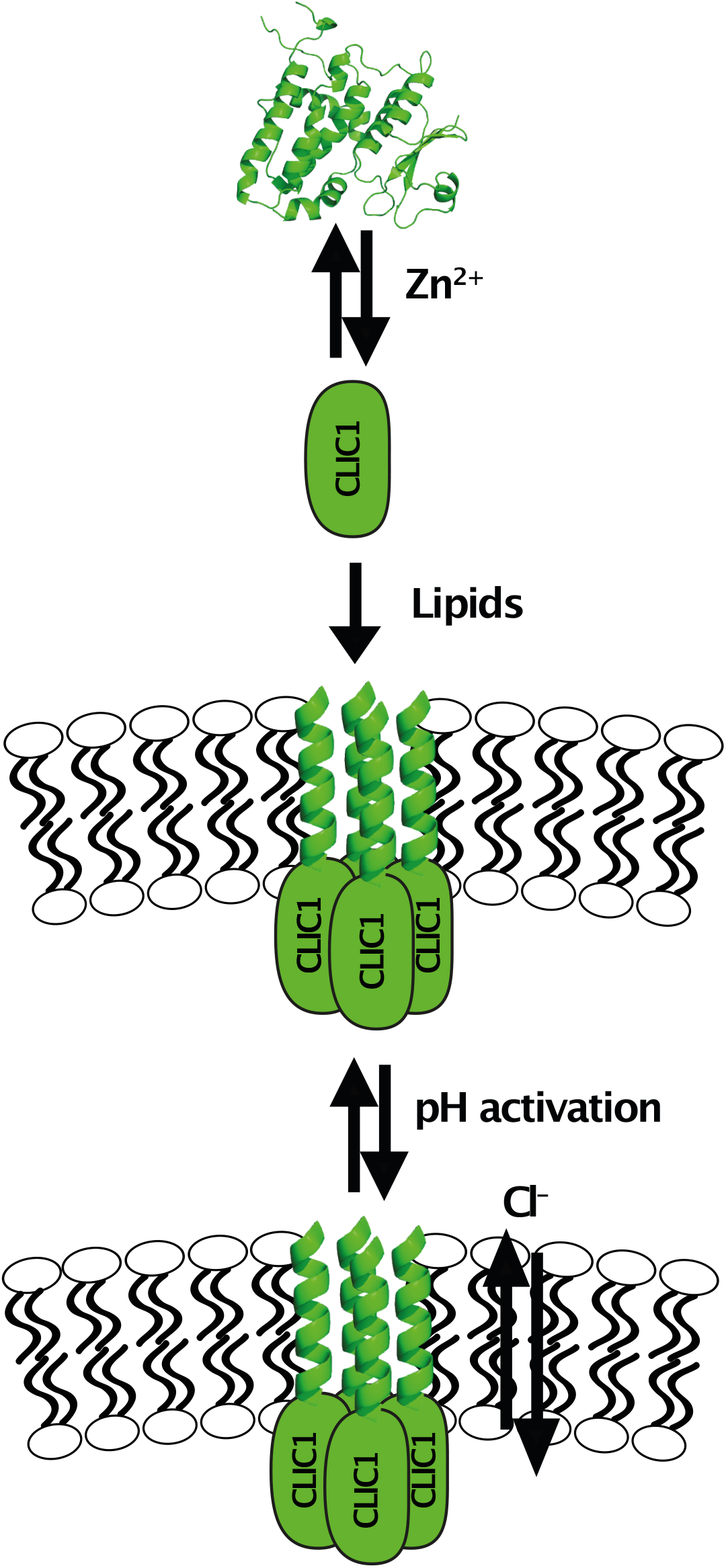
A model for CLIC1 insertion. CLIC1 binds to Zn^2+^, inducing a conformational change that triggers membrane insertion. CLIC1 chloride channel is then activated at low pH.

Immunofluorescence experiments in HUVEC cells revealed a similar level of inhibition of CLIC1 membrane relocalisation with the Zn^2+^ chelator TPEN or the chloride channel inhibitor NPPB. While the mechanism of NPPB inhibition of chloride conductance is still unknown, our results seem to suggest that it blocks Zn^2+^ activation of CLIC1, capturing the protein in the soluble state. Interestingly, despite its non-specific weak inhibition of many families of chloride channels, NPPB is a strong inhibitor of Ca^2+^ activated chloride channels (Verkman and Galietta, 2009), what could suggest a general mechanism preventing divalent cation binding.

Divalent cations have been related to the membrane insertion properties of proteins of the annexin family (Rosengarth and Luecke, 2003), as well as the E1 membrane protein of rubella virus (Dubé et al., 2016) and the amyloidogenic peptide amylin (Sciacca et al., 2013), promoting the interactions of the soluble forms of these proteins with negatively charged lipids. CLIC1 has been shown to have maximal chloride efflux activity in the extract of soybean lipids asolectin, which is rich in the lipid classes Phosphatidylethanolamine (PE), Phosphatidylcholine (PC) and the negatively charged Phosphatidylinositol (PI) (Tulk et al., 2002), supporting the role of divalent cations in CLIC1 membrane insertion. Annexin membrane association is expected to occur due to the exposure of an otherwise buried amphipathic segment upon binding to calcium ions. CLIC1 contains a region (residues 24-41) with moderate hydrophobicity and a moderate hydrophobic moment that could, in a similar mechanism, detach from the protein’s globular structure upon divalent cation binding, become exposed to the solvent and mediate association with the membranes, likely forming a helix. The hydrophobicity of this segment would also explain the aggregation of the protein at higher concentrations of calcium and zinc ions in the absence of lipids. Interestingly, the same region in the structurally homologous Glutathione S-transferase is not hydrophobic, suggesting that this helix plays a different role in the CLIC family.

While the structural rearrangements involved in this process are not yet fully understood, the molecular switch between the soluble and membrane bound forms is now elucidated. This enables, for the first time, the manipulation of CLIC1 localisation in cellular systems and the recombinant production of CLIC1 samples in the chloride channel form in membrane mimetic systems (Medina-Carmona et al., 2020), and provides a clear mechanism for the channel formation process of this unusual and clinically important protein.

## METHODS

### Protein Expression and Purification

The Human CLIC1 gene (clone HsCD00338210 from the Plasmid service at HMS) was cloned into a pASG vector (IBA) containing an N-terminal twin strep tag and a TEV cleavage site, and into a pWaldo vector (Drew et al., 2001) containing a C-terminal GFP and a TEV cleavage site. CLIC1 was expressed recombinantly in the C43 *E*.*coli* strain (Lucigen). The cells were lysed by sonication, and the membrane and soluble fractions were separated by ultracentrifugation at 117734 g. Membrane-bound CLIC1 can be extracted using a mixture of 1% DDM (Glycon) and 1% Triton X-100, resulting in solutions with CLIC1 soluble form. Both fractions were purified separately in the absence of any detergent using affinity chromatography with a Strep-Tactin XT column. The elutions were pooled and cleaved with TEV protease, and subsequently gel filtrated using a Superdex200 Increase column (GE) in either 20 mM HEPES buffer with 20 mM NaCl at pH 7.4 or 20 mM Potassium Phosphate buffer with 20 mM NaCl at pH 7.4.

#### Chloride efflux assays

U87 cells were maintained in DMEM media supplemented with 10% FBS and 1% Penicillin/Streptomycin at 37°C, 95% humidity and 5% CO_2_.

For MQAE (*N*-(Ethoxycarbonylmethyl)-6-Methoxyquinolinium Bromide) (ThermoFisher) assays U87 cells were seeded into dark 96-well microtiter flat-bottom plates at 4 × 10^4^ cells·well^-1^ in 100 µL volumes DMEM media supplemented with 10% FBS and 1% Penicillin/Streptomycin and incubated overnight at 37°C, 95% humidity and 5% CO_2_. Cells were then stained with 8 mM MQAE for 2 h (C et al., 2003), and subsequently washed three times with PBS. Non-stained cells were also included. A variety of conditions were evaluated after MQAE loading. In conditions where inhibitors were used, these – 5 µM TPEN or 10 µM IAA-94 – were added and incubated for 10 min. Finally, 10 µM ionomycin were added and repetitive fluorescence measurements (every 1 minute for 80 minutes) were initiated immediately using an Omega fluorescence plate reader (excitation, 355 nm; emission, 460 nm). Six separate wells were used for each biological repeat. Mean and standard deviations were calculated for each condition and time.

#### Chloride Efflux Assays with purified CLIC1

CLIC1 chloride channel activity was assessed using the chloride selective electrode assay described previously (Tulk et al., 2002). Recombinant CLIC1 was purified as previously described (Tulk et al., 2002) except EDTA was included at 1 mM in all steps following the thrombin digestion until the final dialysis change. Unilamellar asolectin or purified lipid (consisting of 80% palmitolyl oleolyl phophotidyl choline, 10% palmitoyl oleoyl phosphotidyl serine, and 10% cholesterol) vesicles were prepared at 20 mg/mL in 200 mM KCl, 5 mM HEPES (pH 7.4). CLIC1 protein at 1 μM final concentration was incubated in the presence or absence of divalent cation as indicated in 380 μL volume of 200 mM KCl, and 10 mM MES pH 5.5 or HEPES pH 7.5. 20μl of lipid suspension were added to the protein mixture with vortexing and incubated for 5 minutes at room temperature. The lipid mixture was then applied to a 3.5 ml Biogel-P6DG spin column previously equilibrated in 330 mM sucrose. The eluate from the spin column was added to cup containing 330 mM sucrose, 10 μM KCl, 10 mM MES pH 5.5 or HEPES pH 7.5 as appropriate, with continuous monitoring of extravesicular chloride concentration by an ion selective electrode. Valinomycin was added to 1 μM to initiate voltage driven Cl efflux, followed by Triton X-100 to 1% to release remaining intravesicular chloride. The initial rate of fractional chloride efflux after addition of valinomycin is taken as the chloride permeability.

#### Fluorescence Microscopy

Giant unilamellar vesicle formation was carried out using a protocol adapted from (Manley and Gordon, 2008; Veatch, 2007). An Asolectin lipid stock was prepared in 50 mM HEPES, 50 mM NaCl pH 7.4 buffer. 2 μl/cm^2^ of 1 mg/ml lipid mixed with 1 mM Nile red lipophilic stain (ACROS Organic) was applied to two ITO slides and dried under vacuum for 2 hours. 100 mM sucrose, 1 mM HEPES pH 7.4 buffer was used to rehydrate the lipids in the described chamber. 10 Hz frequency sine waves at 1.5 V were applied to the chamber for 2 hours. Liposomes were recovered and diluted into 100 mM glucose, 1 mM HEPES, pH 7.2 buffer. For all four assays 90 nM CLIC1-GFP was incubated with the GUVs with either 0.5 mM ZnCl_2,_ 0.5 mM CaCl_2_, or were left untreated and incubation at room temperature for ten minutes. Microscopy for each assay was performed in an 8 well Lab-Tek Borosilicate Coverglass system (Nun) with a Zeiss LSM-880 confocal microscope using 488 nm and 594 nm lasers. All images were processed with Zen Black software.

For immunofluorescence assays cells were seeded into 24 well plates onto sterile microscopy slides for next day treatment. 24 hours post seeding cells were washed with Tris-buffered saline (TBS), transferred to phosphate free media and left untreated or treated with 10 μM ionomycin or 10 μM *N,N,N*′,*N*′-Tetrakis(2-pyridylmethyl)ethylenediamine(TPEN) as required. Fixation with 4% formaldehyde for 15 minutes was carried out at two hours post treatment. The cells were then washed and permeabilised for 10 minutes with 0.1% Triton in TBS and washed twice with TBS to remove any detergent. Following a TBS wash step the cells were blocked at room temperature with 2% BSA for 1 hour. Primary incubation was carried out overnight at 4 °C with a 1:50 dilution of monoclonal mouse CLIC1 antibody (Santa Cruz Biotechnology, clone 356.1). After primary incubation, 3 wash steps were carried out, prior to 1 hour incubation with secondary antibody at a 1:1000 dilution (Alexa Fluor 488 donkey anti-mouse, Life Technologies). A further 3 washes followed, then nucleus staining with NucBlu Live Cell Stain (Invitrogen) for 20 minutes. The slides were washed a final time and mounted with ProLong Gold Antifade (Invitrogen). All microscopy slides were viewed with a Zeiss LSM-880 confocal microscope using 405 nm, 488 nm, 633 nm lasers. All images were processed with Zen Black and Zen Blue software.

#### Microscale Thermophoresis

A recombinantly expressed C-terminal GFP tag construct of CLIC1 was used. Titrations were performed at a constant protein concentration of 0.8μM in HEPES buffer (20mM HEPES, 20mM NaCl, pH 7.4) in the presence of 80μM asolectin vesicles, with increasing concentration of a solution of Zn_2_SO_4_ in the HEPES buffer and a final volume of 20μL. Prepared samples were filled into Monolith NT.115 capillaries (NanoTemper). Measurements were recorded on a Monolith NT.115 at 30C, excited under blue light, medium MST power and 15% excitation power. The data were analysed using MO Affinity Analysis software (NanoTemper) and fitted using the K_d_ model.

## ACKNOWLEDGEMENTS

We thank Dr N. Fili and Dr. J. Rossman for help with confocal imaging, and Dr G.S. Thompson and Dr. D.A.I. Mavridou for feedback on the manuscript. RME is supported by a fellowship from the Alfonso Martin Escudero Foundation. VAS is supported by Barts Charity’s Rising Star programme. We acknowledge support from the Wellcome Trust Seed Award (207743/Z/17/Z).

## CONTRIBUTIONS

JLOR, LV and ACH designed experiments. LV performed protein expression and purification, preliminary biophysical assays and chloride efflux measurements. ACH performed fluorescence assays, GUV experiments and cell imaging. JC measured the MST binding experiments. RME collected the chloride efflux assays in cells. VAS performed fluorescence imaging experiments in HUVEC cells. DC aided with cell and GUV imaging experiments at Kent. JE performed the chloride efflux studies in purified protein. JLOR, LV and ACH prepared figures. JLOR supervised the project and prepared the manuscript. JLOR, LV and ACH edited the manuscript. LV and ACH contributed equally to the work.

## FIGURES

## SUPPLEMENTAL FIGURES

**Figure S1-** Fluorescent microscopy images of Asolectin GUVs labelled with Nile red dye incubated with GFP-labelled CLIC1 in the absence of divalent cations and in the presence of 500 μM of Zn^2+^ or Ca^2+^.

