## Supplementary figures and images for "Zn^2+^ triggered two-step mechanism of CLIC1 membrane insertion and activation into chloride channels"

### Supplemental Figure 1

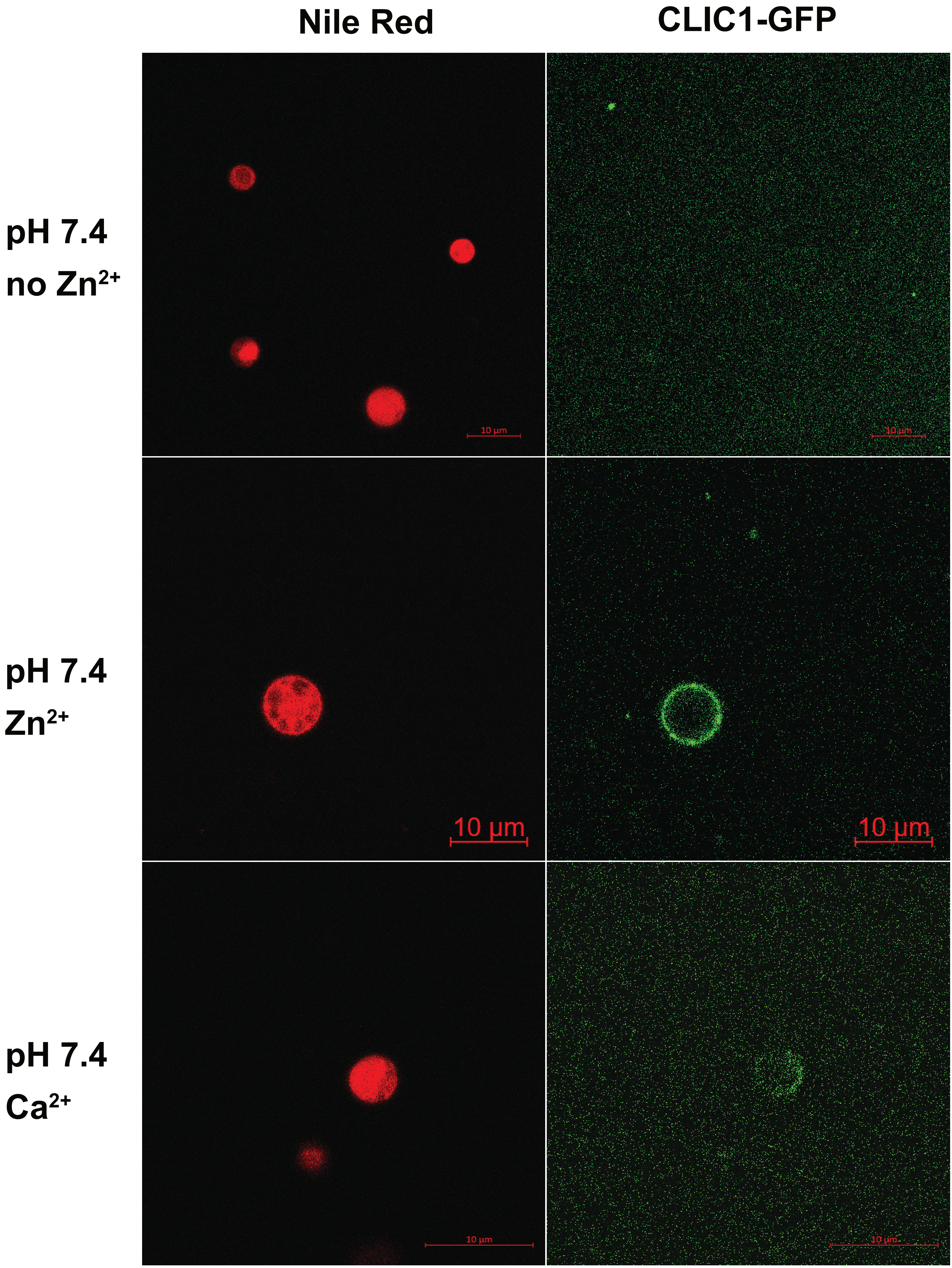
